# Chromatin accessibility changes induced by the microbial metabolite butyrate reveal possible mechanisms of anti-cancer effects

**DOI:** 10.1101/2021.03.30.437582

**Authors:** Matthew G. Durrant, Brayon J. Fremin, Abhiram Rao, Emily Cribas, Stephen Montgomery, Ami S. Bhatt

## Abstract

Butyrate is a four-carbon fatty acid produced in large quantities by bacteria found in the human gut. It is the major source of colonic epithelial cell energy, can bind to and agonize short-chain fatty acid G-protein coupled receptors and functions as a histone deacetylase (HDAC) inhibitor. Anti-cancer effects of butyrate are attributed to a global increase in histone acetylation in colon cancer cells; however, the role that corresponding chromatin remodeling plays in this effect is not fully understood. We used longitudinal paired ATAC-seq and RNA-seq on HCT-116 colon cancer cells to determine how butyrate-related chromatin changes functionally associate with cancer. We detected distinct temporal changes in chromatin accessibility in response to butyrate with less accessible regions enriched in transcription factor binding motifs and distal enhancers. These regions significantly overlapped with regions maintained by the SWI/SNF chromatin remodeler, and were further enriched amongst chromatin regions that are associated with ARID1A/B synthetic lethality. Finally, we found that butyrate-induced chromatin regions were enriched for both colorectal cancer GWAS loci and somatic mutations in cancer. These results demonstrate the convergence of both somatic mutations and GWAS risk variants for colon cancer within butyrate-responsive chromatin regions, providing a molecular map of the mechanisms by which this microbial metabolite might confer anti-cancer properties.

**Highlights:**

- Chromatin accessibility changes longitudinally upon butyrate exposure in colon cancer cells.
- Chromatin regions that close in response to butyrate are enriched among distal enhancers.
- There is strong overlap between butyrate-induced peaks and peaks associated with SWI/SNF synthetic lethality.
- Butyrate-induced peaks are enriched for colorectal cancer GWAS loci and somatic variation in colorectal cancer.

## Introduction

Dietary components that reach the colon are used by the colonic microbial community and yield diverse metabolites. Among these are fermentation products known as short-chain fatty acids (SCFAs) (Wu et al., 2018). Butyrate is among the most well-studied SCFAs in the context of colorectal cancer (Donohoe et al., 2012). Butyrate has a variety of functions, including HDAC inhibition (Donohoe et al., 2012) and binding to GPCR receptors (Husted et al., 2017). Colonic epithelial cells metabolize butyrate as a primary source of energy, but due to the Warburg effect, glucose is utilized instead of butyrate as the primary energy source in colon cancer cells (Donohoe et al., 2012; Fleming et al., 1991; Roediger, 1982). It is hypothesized that this allows butyrate to accumulate intracellularly and act as a potent HDAC inhibitor in colon cancer cells. This accumulation of butyrate further manifests in global increases in histone acetylation and subsequent chromatin remodelling that are expected to underlie its anti-cancer effects on colon cancer cells, including diminished proliferation (Donohoe et al., 2012). Such chromatin accessibility changes in response to butyrate have been previously studied in rumen epithelial cells (Fang et al., 2019) and leukemia cells (Frank et al., 2016). However, the specific changes in chromatin accessibility and associated gene expression changes induced by butyrate exposure in colon cancer cells have not been well characterized.

HDAC inhibition has been linked to a number of protein complexes involved in cancer, including the SWI/SNF (SWItch/Sucrose Non-Fermentable) complex (Fukumoto et al., 2018). SWI/SNF complex subunits are collectively mutated in approximately 20 percent of all cancers (Garraway and Lander, 2013; Kadoch et al., 2013; Mathur et al., 2017). ARID1A is the most frequently mutated subunit in this complex. ARID1A mutations sensitize cancer cells to HDAC inhibition (Fukumoto et al., 2018). ARID1A loss has also been shown to drive colon cancer in mice via impairment of enhancer-mediated gene regulation (Mathur et al., 2017). However, combinations of loss of function in SWI/SNF complex subunits can induce synthetic lethality in cancer cells. For example, a loss of function of both ARID1A and ARID1B induces synthetic lethality in HCT-116 colon cancer cells (Kelso et al., 2017). Though HDAC inhibition and SWI/SNF mutations and regulation are linked in the context of cancer, their mechanisms of interaction and the role of butyrate remain unclear.

The interactions between the human gut microbiome and common germline genetic variants and somatic mutations in the host is an area of active research providing the potential for discovery of new cancer risk factors and treatments. One recent study demonstrated that the gut microbe metabolite gallic acid may interact with somatic mutations in p53 to influence oncogenesis (Kadosh et al., 2020). Butyrate is considered to be an ideal candidate to discover such gene-environment interactions due to its diverse cellular functions and direct relationship to dietary fiber intake (Bultman, 2014). In this study, we aim to identify how butyrate modulates the effect of both common germline variants and somatic mutations that influence colorectal cancer through butyrate-stimulated chromatin accessibility changes in human host cells.

## Results

### Butyrate decreases chromatin accessibility in distal enhancer regions

The HDAC-inhibitory effect of butyrate is well-documented (Donohoe et al., 2012). HDAC inhibition suggests greater histone acetylation throughout the genome, which our own experiments confirmed (Fig. S1). To test the effect of butyrate on the chromatin conformation of colon cancer cells, we exposed HCT-116 cells to control conditions or butyrate. We generated longitudinal ATAC-Seq libraries for three time points at 9, 18, and 24 hours for the butyrate-exposed samples and the controls. We sequenced a total of 746,181,642 ATAC-Seq reads (range = 44,659,678-138,802,186 reads per replicate). For each time point, we observed strong nucleosome phasing and transcription start site enrichment (Fig. S2). Differential accessibility analysis indicated the number of peaks opening and closing in response to butyrate treatment was roughly equal over the time course (Fig. 1A & Table S1). In total, 6,128 peaks were found to be differentially accessible during at least one time point (*FDR* < 0.1, |log2(Fold Change)*l* > 1; Fig. 1A), representing ~12% of the 52,530 peaks tested (Table S2). Principal components analysis demonstrated that butyrate treatment was the primary source of variation (Fig. S3A). Furthermore, we observed that the total number of differentially accessible peaks increased as time progressed and subsequently referred to chromatin regions that became less accessible in response to butyrate as “closed peaks” and regions that became more accessible as “open peaks.” While the opening of chromatin was expected given the function of butyrate as an HDAC inhibitor, the large number of closed peaks despite global increases in histone acetylation was surprising, albeit not unprecedented (Frank et al., 2016).

**Figure 1.**
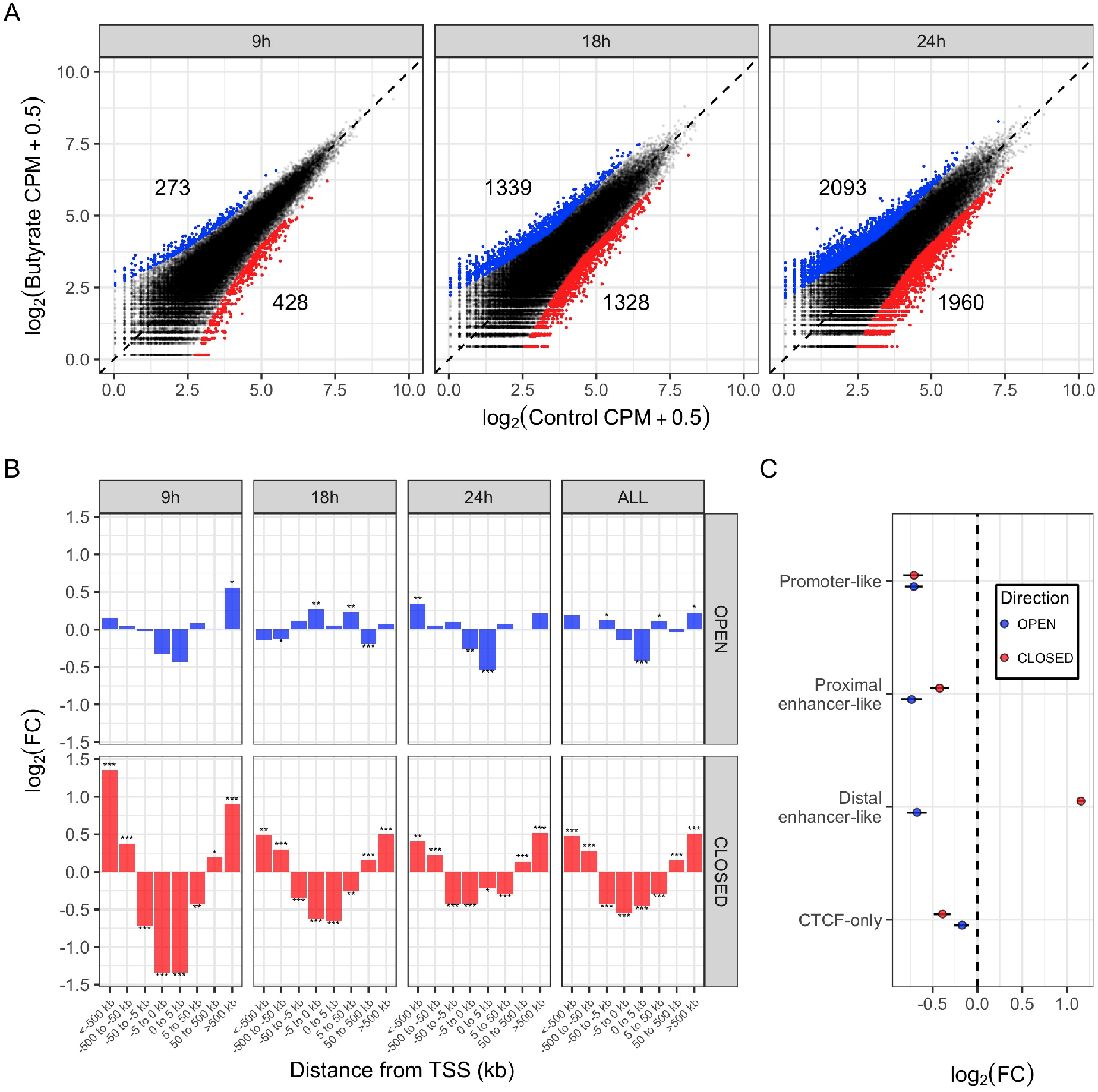
Butyrate decreases chromatin accessibility in distal enhancer regions. (A) Differentially accessible genomic regions as measured by ATAC-seq. Showing three time points following butyrate treatment compared to untreated controls. The average log2(CPM) is shown. Blue points are regions that become significantly more accessible in response to butyrate treatment (open peaks; FDR < 0.05 and log2(FC) > 1). Red points are regions that become significantly less accessible in response to butyrate treatment (closed peaks; FDR < 0.05 and log2(FC) < −1). Numbers above the blue points and below the red points indicate the total number of significantly more open and closed peaks at each time point, respectively. (B) Enrichment of open and closed regions at different distances from the nearby transcription start sites (TSS). Enrichment was calculated using a hypergeometric test, with the log2(Hypergeometric Fold Change) values shown on the y-axis. Positive values along the y-axis indicate that the regions in the given distance bin are enriched relative to the background of all tested regions, negative values indicate that they are depleted. The column labeled “ALL” indicates all open/closed peaks across all time points considered together. ‘*’ indicates FDR < 0.05, ‘**’ indicates FDR < 0.01, ‘***’ indicates FDR < 0.001. Negative numbers along the x-axis indicate regions that are upstream of a nearby TSS, and positive numbers are downstream of a nearby TSS. (C) Enrichment of open and closed peaks overlapping with different HCT116 candidate cis-regulatory elements (CREs) as determined from ENCODE data and made available through the SCREEN web interface. Only shown are enrichment/depletion of all significantly open and closed peaks aggregated across all three time points. Lines through each point indicate the 95% bootstrapped confidence interval of the log2(Hypergeometric Fold Change).

Longitudinal gene expression data was also generated using RNA-seq. We sequenced a total of 810,869,958 RNA-Seq reads (range = 65,806,356-107,642,896 reads per replicate). Differential expression analysis indicated that approximately 78.4% of genes were differentially expressed in response to butyrate during at least one time point (FDR < 0.05), and 69.7% of those genes were differentially expressed above |log_2_(Fold Change)| > 1. Principal components analysis of RNA-seq data suggested that butyrate treatment and time after exposure were the primary sources of variation in the data (Fig. S3B). Gene set enrichment analysis (GSEA) indicated several pathways that were differentially expressed; for example, we observed a significant down-regulation of E2F targets and G2M checkpoint genes, indicating that butyrate strongly impacted cell growth (Fig. S3C). By combining chromatin-accessibility and gene expression data, we also found evidence that differentially-accessible regions were associated with differentially-expressed genes (Fig. S3D).

Next, we inspected the distribution of differentially accessible peaks across the genome. We found that closed peaks were particularly enriched in intergenic regions that were distal to the nearest TSS (Fig. 1B). Peaks that were distantly upstream (<-500 kbp from TSS) or downstream (>500 kbp from TSS) were found to be the most strongly enriched in closed peaks across all time points, and especially at 9 hours following butyrate exposure. By contrast, the genomic pattern observed in the open peaks was much less conserved across all three time points, and the enrichment/depletion effect sizes were relatively modest. This suggests that the effect of butyrate on closing peaks was more targeted and consistent than the effect on the opening peaks.

To determine if butyrate-induced peaks were enriched/depleted in cis-regulatory elements (CREs), we used ENCODE data made available through the SCREEN web interface to identify candidate CREs (Fig. 1C). Both closed and open peaks were depleted of promoter-like, proximal enhancer-like, and CTCF-only CREs. Closed peaks were strongly enriched (log2(FC) > 1) for distal enhancer-like elements. Taken together, these data indicate that butyrate induced both the closing and opening of peaks across the genome, but the closed peaks were particularly enriched for distal enhancer regions, especially at 9 hours after butyrate exposure, while the genomic location of the open peaks appeared more sporadic across the time points.

### Butyrate-induced closed peaks are enriched for transcription factor binding, including SWI/SNF complex, AP-1 complex, and TEAD binding sites

We next investigated whether differential peaks were significantly enriched for specific transcription factor binding targets. We compared our differentially accessible peaks to previously generated ChIP-seq peaks for the HCT-116 cell line (Fig. 2A). Open peaks were strongly depleted for most of the ChIP-seq signals, especially at 18 and 24 hours after butyrate exposure. Closed peaks, in contrast, showed significant enrichment in ChIP-seq signals across all three time points. In particular, SWI/SNF subunits SMARCA4 and SMARCC1 ChiP-seq peaks were the most strongly enriched among the butyrate-induced closed peaks at 9 hours, with a log2(FC) of 2.46, suggesting that butyrate-induced closure of SWI/SNF binding sites is a particularly strong signal, especially early on following butyrate exposure. Binding sites for AP-1 complex subunits FOSL1 and JUND were also strongly enriched in closed peaks, as well as TEAD, CEBP, CBX3, SP1, SRF, JAK2, and ATF3 binding sites.

**Figure 2.**
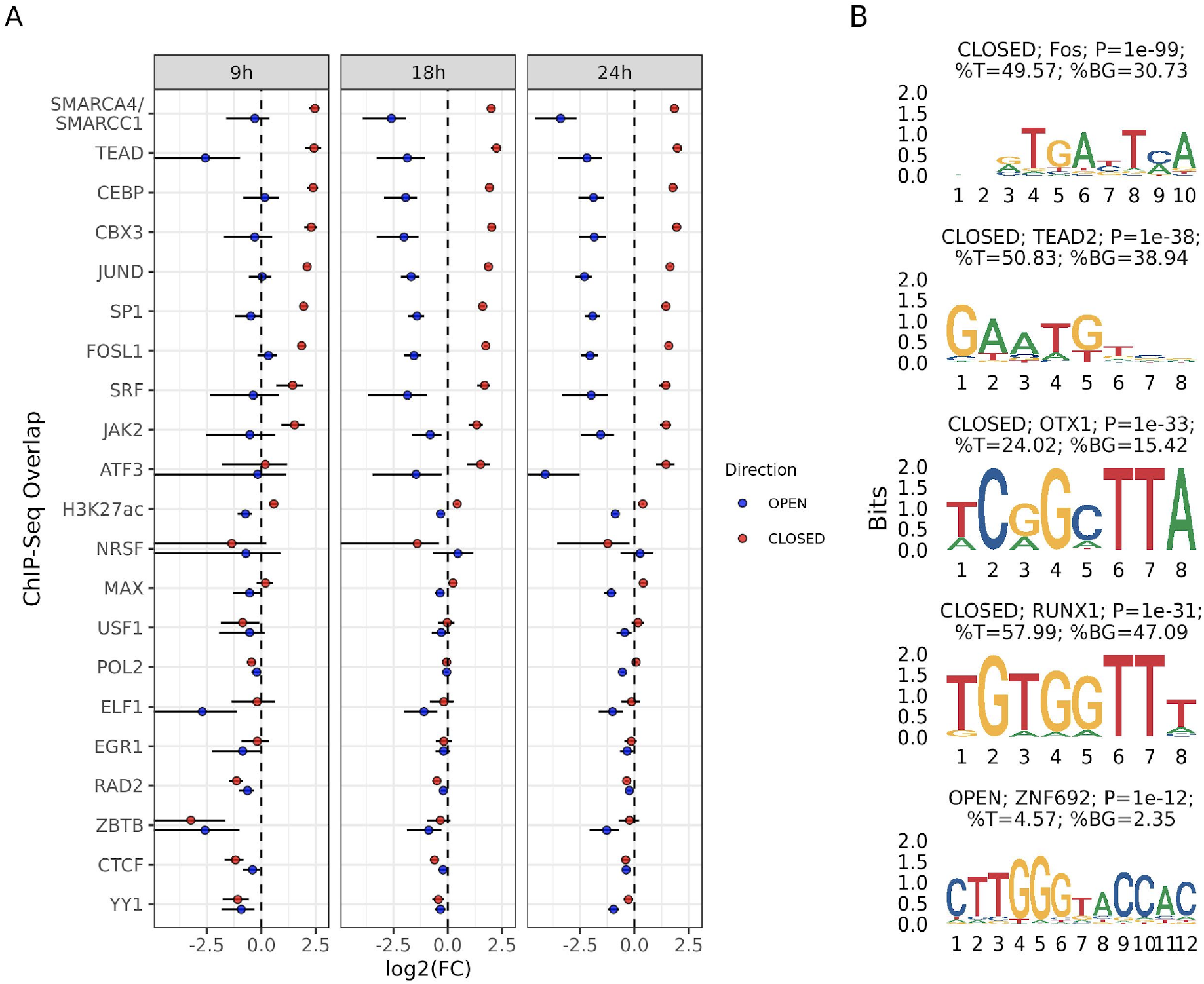
Butyrate-induced closed peaks are enriched for transcription factor binding, including SWI/SNF complex, AP-1 complex, and TEAD binding sites. (A) Enrichment of Butyrate-induced peaks with ChIP-seq peaks. Each row corresponds to a different ChIP-Seq experiment performed on HCT-116 cells. Points indicate open (blue) and closed (red) peaks at 9, 18, and 24 hours after butyrate treatment. Lines through each point indicate the 95% bootstrapped confidence interval of the log2(Hypergeometric Fold Change). (B) Top *de novo* motifs enriched in all significant butyrate-induced peaks as identified by the HOMER motif finding software. Showing the top 4 *de novo* motifs found across all closed peaks, and the only *de novo* motif in the open peaks that meets the HOMER-recommended significance threshold. Titles of each motif indicate if they were enriched in open/closed peaks, the protein with the best-matching known motif, the P-value of the enrichment statistic, the percentage of target (%T) sequences that contain the motif, and the percentage of background (%BG) sequences that contain the motif.

We also used the HOMER motif finding software to identify enriched motifs *de novo* in both closed and open peaks (Fig. 2B) (Heinz et al., 2010). The most enriched motif in closed peaks was similar to a Fos-associated binding motif, where 49.57% of all closed peaks contained such a motif, compared to 30.73% of the background regions. Other enriched motifs in closed peaks included those associated with the TEAD2, OTX1, and RUNX1 transcription factors. The open peaks contained only one significant *de novo* motif (using the HOMER-recommended significance cutoff) associated with transcription factor Zinc Finger Protein 692 (ZNF692). Taken together, these data suggest that butyrate exposure results in the selective closure of multiple distal regulatory elements and chromatin loops that are being actively maintained by the AP-1 complex and the SWI/SNF complex.

### Butyrate-induced peaks significantly overlap with regions associated with synthetic lethality of SWI/SNF complex subunits ARID1A/B

We found that the chromatin accessibility changes that we observed due to butyrate exposure were similar to those reported in a study conducted by Kelso et al. (Kelso et al., 2017). In this study, Kelso et al. investigated the chromatin accessibility changes that occurred in the HCT-116 cell line in response to gene deletion and gene knockdown of two important SWI/SNF complex subunits, ARID1A and ARID1B. The SWI/SNF complex maintains chromatin architecture, and mutations in the ARID1A subunit are commonly found in cancer (Kadoch et al., 2013). Deficiency of the ARID1B subunit is synthetically lethal with ARID1A mutation, and it was this synthetic lethality that Kelso et al. further investigated in their study.

We analyzed the Kelso et al. (2017) publicly available ATAC-seq data using the same pipeline as we used for our own butyrate-treated ATAC-seq data (Fig. 3A, see Methods). Using the 52,530 peaks identified in our study, we identified differentially accessible peaks in the three treatments relative to our untreated control, as well as the ARID1A -/- & ARID1B KD treatment relative to the ARID1A -/- treatment as a control. The ARID1A -/- & ARID1B KD vs. ARID1A -/- comparison identified peaks that were specific to the synthetic lethality phenotype. We identified 12,324 total differentially accessible peaks, with 1,908, 5,080, and 141 peaks that opened in the ARID1A -/-, ARID1A -/- & ARID1B KD, and ARID1A -/- & ARID1B KD vs. ARID1A -/- treatments, respectively, and 3,748, 6,250, and 1,072 peaks that closed in the same three treatments, respectively.

**Figure 3:**
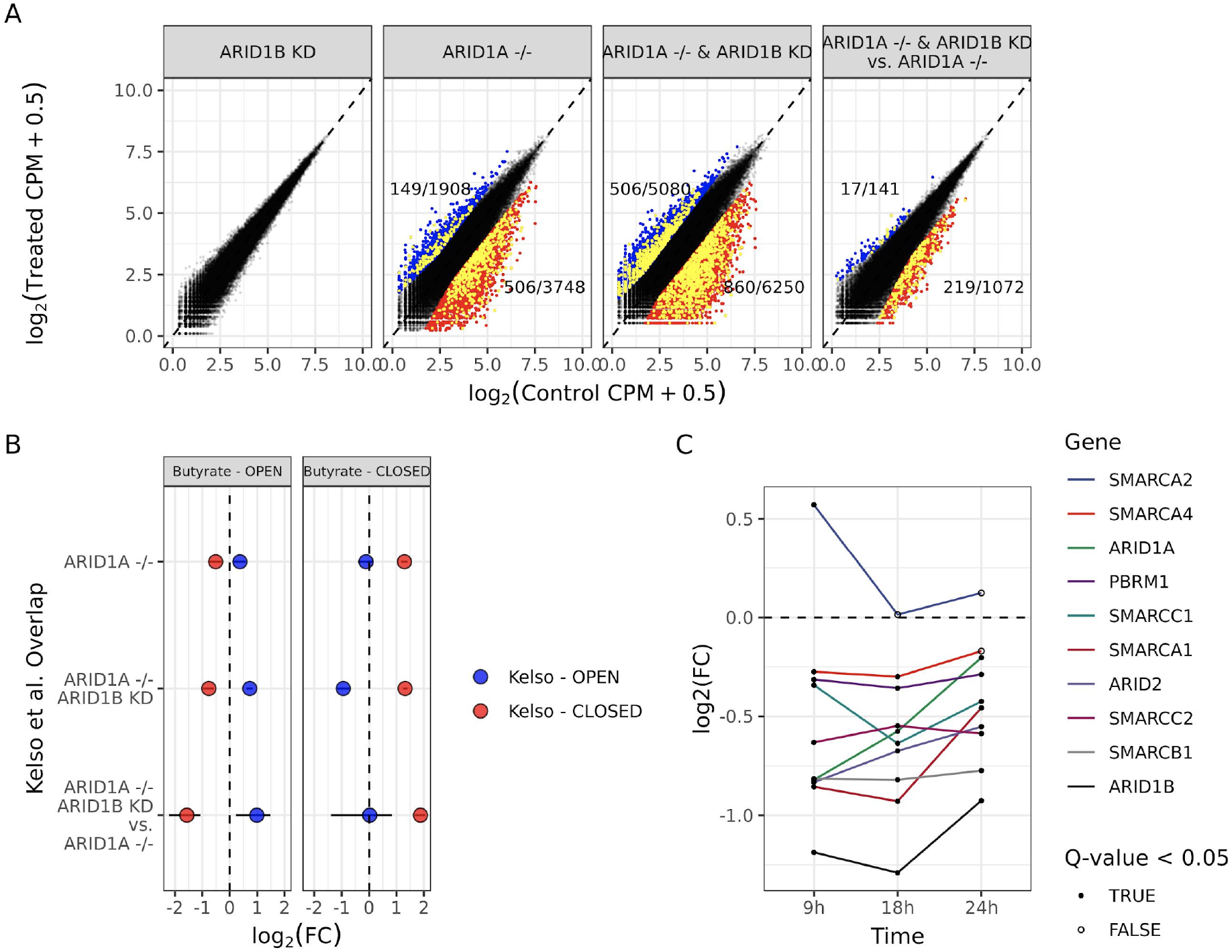
Butyrate-induced peaks significantly overlap with regions associated with synthetic lethality of SWI/SNF complex subunits ARID1A/B. (A) Differentially accessible genomic regions as measured by ATAC-seq of HCT-116 from data presented in the Kelso et al. (2017) study. Showing differential peaks under three conditions: shRNA knock down of ARID1B (ARID1B KD), homozygous loss of ARID1A (ARID1A -/-), and both conditions simultaneously (ARID1A -/- & ARID1B KD). Differential ATAC-seq peaks of these three conditions were determined relative to untreated controls, and were also determined in the ARID1A -/- & ARID1B KD treatment relative to the ARID1A -/- control (ARID1A -/- & ARID1B KD vs. ARID1A -/-). Blue points are regions that become significantly more accessible in response to each treatment (open peaks; FDR < 0.05 and log2(FC) > 1). Red points are regions that become significantly less accessible in response to each treatment (closed peaks; FDR < 0.05 and log2(FC) < −1). Yellow points are also differentially accessible in at least one of the butyrate-treated time points in this study. Fractions above the blue points and below the red points indicate the total number of significant open and closed peaks at each time point in the denominator, respectively, and the total number that overlap with butyrate-induced differentially accessible peaks in the numerator. (B) Enrichment of butyrate-induced open and closed peaks among the differentially accessible peaks in the Kelso et al. (2017) study. Blue and red points indicate the log2(Hypergeometric Fold Change) of butyrate-induced open and closed peaks in the three treatment conditions. Lines through each point indicate the 95% bootstrapped confidence interval of the log2(Hypergeometric Fold Change). (C) The log2(Fold Change) in gene expression of 10 subunits of the SWI/SNF complex as measured by RNA-seq relative to untreated controls. Showing time points 9 hours, 18 hours, and 24 hours after butyrate treatment. ARID1B is the only subunit that is significantly down-regulated in response to butyrate treatment, but the direction of the effect is consistent in the direction of down-regulation across all 3 time points for 9 of the 10 subunits.

We found that across the Kelso et al. (2017) differentially accessible peaks, there was significant overlap with butyrate-induced differentially accessible peaks (Fig. 3B). Butyrate-induced open peaks significantly overlapped with the Kelso et al. open peaks in all treatment conditions, and they were significantly depleted among the Kelso et al. closed peaks. Butyrate-induced closed peaks were significantly enriched among the Kelso et al. closed peaks in all treatment conditions, with the strongest effect among the peaks associated with synthetic lethality (ARID1A -/- & ARID1B KD vs. ARID1A -/- peaks; P < 0.001; log2(Hypergeometric Fold Change) = 1.87). Among the peaks associated with ARID1A/B synthetic lethality, 19.5% were also differentially accessible in the same direction in at least one of the butyrate-treated time points.

To test if the SWI/SNF subunit may be disrupted in response to butyrate treatment, we analyzed SWI/SNF subunit gene expression using RNA-seq (Fig. 3C). We found that all subunits of the SWI/SNF complex were significantly down-regulated during at least one time point (FDR < 0.1), with the exception of SMARCA2 which was significantly upregulated at 9 hours after butyrate exposure. The most significantly down-regulated gene belonging to the SWI/SNF complex across all three timepoints was ARID1B. Taken together, this indicates that the effect that butyrate has on these regions may be due in part to genetic down-regulation of a large number of the SWI/SNF complex subunits.

### Differentially accessible chromatin regions are enriched for colorectal cancer GMAS loci and cancer-associated somatic mutation

To assess the role that butyrate-induced differential accessible regions may have to cancer, we first assessed if these regions were enriched for colorectal cancer heritability. Stratified LD-score regression has been used to determine if regions surrounding genes expressed in tissue-specific manner are enriched for disease heritability as measured by GWAS summary statistics (Finucane et al., 2018). Given the known association between butyrate and colorectal cancer, we used this same approach to determine if butyrate-responsive peaks are associated with colorectal cancer heritability.

The results of our heritability enrichment analysis indicated that open peaks were significantly enriched for colorectal cancer heritability (P = 0.019), while closed peaks were not (P = 0.790), where positive normalized effect sizes indicate heritability enrichment (Fig. 4A). When we restricted our analysis to only distal peaks where the nearest gene is greater than 50 kilobases away, we observed that the enrichment for colorectal cancer heritability slightly increases in the open peaks (P = 0.004), and the closed peaks remain non-significant.

**Figure 4:**
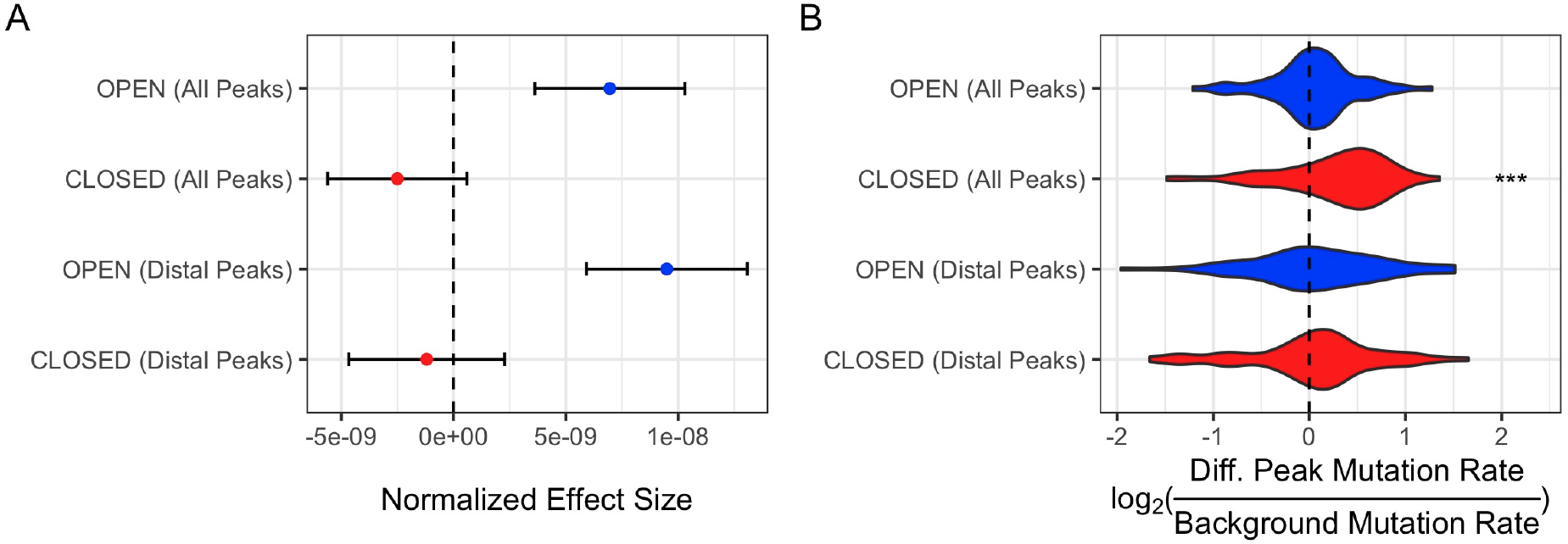
Differentially accessible chromatin regions are enriched for colorectal cancer GWAS loci and cancer-associated somatic mutation. (A) Heritability enrichment statistics for colorectal cancer as calculated using stratified LD-score regression and colorectal cancer GWAS summary statistics. Showing the normalized effect size of enrichment ± standard error. Blue points refer to open peaks, red points refer to closed peaks. (B) Somatic mutation enrichment in butyrate-responsive peaks in 60 colorectal cancer samples. Showing the distribution of log2(Diff. Peak mutation Rate / Background Mutation Rate) values on each line. P < 0.001 is represented as three asterisks. To test the “OPEN (Distal Peaks)” peak set, only 57 samples were used because in three samples the relative mutation rate could not be calculated due to low number of mutations.

While this analysis measures colorectal cancer heritability enrichment within butyrate-responsive peaks, we next wanted to investigate the relevance of these peaks to somatic mutation in cancer. We used somatic mutation data generated by the Pan-Cancer Analysis of Whole Genomes (PCAWG), which includes 828 samples from the same number of donors across 16 body sites. We tested sets of peaks to determine if they were enriched or depleted for somatic mutations by comparing their somatic mutation rate to the somatic mutation rate in peaks that were non-responsive to butyrate.

Given that the colon is the site of highest butyrate concentration within the body, we first limited our analysis to the 60 colorectal cancers available (Fig. 4B). In this analysis, we found that closed peaks were enriched for high somatic mutation rates (One-sample t-test; P = 0.0008, Average log_2_(Diff. Peak Mutation Rate / Background Mutation Rate) = 0.258), while open peaks were not (One-sample t-test; P = 0.438; Average log2(Diff. Peak Mutation Rate / Background Mutation Rate) = 0.0438). When limiting to only distal peaks, this enrichment disappeared. As butyrate can translocate into the bloodstream and thus can reach distal body sites, we next repeated this analysis across several different cancer types (Fig. S4). Notably, it was only in colorectal cancer where we observed somatic mutation enrichment in closed peaks. In cancers at various body sites, closed peaks were often significantly depleted of somatic mutations (Bladder, Brain, Breast, Head and Neck, Kidney, Mesenchymal, Ovary, and Skin), and in several body sites open peaks were enriched (Head and neck, Kidney, Lung, Prostate, and Skin).

## Discussion

While many studies have demonstrated strong associations between the gut microbiome composition and various diseases, studying host-microbe interactions has been challenging from a mechanistic perspective (Bhutia et al., 2017). Certainly, alteration of the gut microbiome, in some cases with extreme therapies such as fecal microbiota transplantation, has produced therapeutic benefits in selected circumstances. However, choosing donors can be difficult, and the composition of bacteria is not guaranteed to remain entirely consistent or perform the same roles in the new context (Andremont, 2017; van Beurden et al., 2017; Olesen et al., 2016). A more conventional and controllable approach is to understand underlying mechanisms by identifying the effects that specific microbes and their metabolites elicit on host cells. Butyrate is among the most well studied microbial metabolites, and while its role in modifying the cellular composition of the intestinal lamina propria is known, the impact of butyrate on colonic epithelial cells at the genomic and gene-level is less well understood. In this work, we studied the effects of butyrate, a microbial metabolite, on HCT-116 colorectal cancer cells over time to reveal chromatin accessibility and gene expression changes and their relationship to cancer-related loci.

To characterize the mechanistic links between the microbial metabolite butyrate and specific chromatin accessibility and gene expression changes, we performed paired ATAC-seq and RNA-seq on HCT-116 cells to monitor the effects of butyrate on colon cancer cells over time. We hypothesized that butyrate would increase chromatin accessibility of regions across the genome, driving gene expression changes. Despite global increases in histone acetylation via butyrate, widespread, targeted ‘closing’ of regions of chromatin was more strongly associated with significant effects on the cells. Interestingly, SWI/SNF subunits were collectively downregulated upon butyrate exposure, and the closed chromatin regions we identified were actively maintained by the SWI/SNF complex. This indicates that butyrate alters chromatin accessibility in multiple ways, both directly via HDAC inhibition, and indirectly via downregulation of the SWI/SNF complex, thus disrupting maintenance of chromatin structure.

Additionally, we find that butyrate may influence colorectal cancer susceptibility both in terms of germline variation and somatic mutation, our assumption being that through changing the accessibility of the relevant genomic variants butyrate modulates their downstream phenotypic effects. Interestingly, the open peaks were enriched for germline variation associated with colorectal cancer, while the closed peaks were enriched for somatic variants in colorectal cancer. The specificity of the somatic mutation enrichment to colorectal cancer was further notable, as it suggests that the tissue that is most directly exposed to butyrate is the most relevant in terms of potential gene-environment interactions.

We acknowledge that our conclusions are limited in their scope. Further experiments are necessary to determine the direct mechanism by which butyrate affects SWI/SNF-associated regions. The fact that the SWI/SNF effect is strongest at 9 hours after exposure indicates that this is the strongest initial effect of butyrate exposure, and other transcriptional and chromatin accessibility effects may be downstream consequences of early SWI/SNF inhibition. The associations with germline and somatic colorectal cancer risk also warrants further experimental investigation. The fact that peaks that open in response to butyrate are enriched for heritability as measured by a GWAS of common variants while closed peaks are enriched for somatic variation in cancer is an interesting finding that we were not able to address in the scope of this study. Additionally, it is not clear from this study to what extent these findings can be generalized to non-cancerous cells, where butyrate does not accumulate at high concentrations. Finally, it is not clear to what extent cancerous cells may adapt to high levels of butyrate over time, potentially circumventing the anti-cancer effects of butyrate exposure.

In conclusion, we present evidence that highlights potential mechanisms by which butyrate, a prevalent microbial metabolite, influences colorectal cancer risk. The global effects of butyrate on chromatin accessibility have been observed in the past, and it is likely that other microbial metabolites have similarly dramatic effects on host gene expression and chromatin accessibility. Dietary composition is known to play a role in the production of butyrate, and increasing the production of butyrate in the gut has been proposed as a therapeutic strategy to treat a wide range of human diseases (Canani et al., 2011). We believe that this study can help direct further efforts to develop such therapies and to more thoroughly understand their mechanism of action.

## Methods

### Cell Culture

HCT-116 cells were purchased from Sigma (91091005). HCT-116 cells were grown in Dulbecco’s Modified Eagle’s Medium (DMEM, Sigma-Aldrich) supplemented with 10 percent Fetal Bovine Serum (FBS, Gibco) in T25 flasks. At 60 percent confluency, media was replaced with serum-free DMEM media containing differential quantities of butyrate (Alfa Aesar) for various times. Replicates were exposed to the same conditions in different flasks. At designated times, 50,000 cells from each replicate were frozen in 10 percent Dimethyl sulfoxide (DMSO) and 90 percent FBS to be used for ATAC-Sequencing. The remaining cells in the flask were snap frozen to be utilized for RNA-Sequencing.

### ATAC-Seq

ATAC-seq was performed on 50,000 HCT-116 cells from each treatment. Each treatment was done in biological replicate. All conditions were performed in the same batch. 50,000 cells were established as yielding the highest quality libraries for HCT-116 cells. Protocols for ATAC-Seq performed as described (Buenrostro et al., 2013, 2015). These libraries were pooled and sequenced on a Next-Seq 500 (Illumina), obtaining 101 base pair paired-end data.

### RNA-Sequencing

RNA was extracted from HCT-116 cells using the RNA-Easy Mini Plus Kit (Qiagen). Likewise, each treatment was performed in biological duplicate consistent with the ATAC-seq duplicates. All conditions were performed in the same batch. 2 ug of total RNA, quantified using the Qubit RNA HS kit, was used as input for Tru-seq mRNA Stranded kit (Illumina). Standard Illumina protocols were performed. The libraries were pooled and sequenced on a Next-seq 500 (Illumina), obtaining 101 base pair paired-end data.

### ATAC-seq Analysis

Sequences were run through the Big Data Script ATAC-seq pipeline created by the Kundaje lab (https://github.com/kundajelab/atac_dnase_pipelines). Data was processed as previously described (Corces et al., 2016; Miyamoto et al., 2018). The Several dependencies were utilized (Daley and Smith, 2013; Langmead and Salzberg, 2012; Quinlan and Hall, 2010). This pipeline utilizes macs2 (Zhang et al., 2008) for peak calling. Peaks were called by merging all of the optimal IDR peak calls from each time point. Read counts per peak were calculated using the bedtools coverage command line utility (Quinlan and Hall, 2010). Prior to calling differential peaks, counts were quantile normalized using the preprocessCore package in R. This was necessary to overcome a strong increase in noise observed in time points 18 and 24 following butyrate treatment. Quantile-normalized counts were then used to identify differentially accessible ATAC-seq peaks using edgeR (Robinson et al., 2010). The two replicates at each time point were individually compared to three control samples taken at 9 hours, 18 hours, and 24 hours. Peaks with a q-value less than 0.1 and a | log2(Fold Change) | > 1 were considered to be differentially accessible. The same workflow was used to analyze the ATAC-seq data produced by Kelso et al, with the exception of using the merged peaks from our study rather than re-calling peaks on their data.

### RNA-sequencing Analysis

Reads were deduplicated using Super Deduper (Petersen et al., 2015) and adapters were trimmed using Trim Galore version 0.6.6 and Cutadapt version 1.18 (Martin, 2011). GRCh37 was used as the reference genome. Reads were aligned using STAR version 2.7.6a (Dobin et al., 2013). These files were sorted and used as input for HTSeq version 0.11.3 (Anders et al., 2015). Raw counts were analyzed using DESeq2 (Love, Huber, and Anders 2014). Genes were only considered if their counts per million (CPM) exceeded 1 in at least one of the replicates of each treatment, and then only if cutoff was met in all treatments, resulting in 13,398 genes. Significance was assigned with a q-value < 0.05 after Benjamini and Hochberg correction (Dabney et al., 2011). Gene set enrichment analysis of differentially expressed genes were performed using Enrichr (Kuleshov et al., 2016).

### Enrichment analysis - distance from transcription start site (TSS)

The distance of each ATAC-seq peak to the nearest gene was calculated by using the GREAT web service version 3.0.0 (McLean et al., 2010). Peaks were associated with nearby genes using the “Basal+extension” approach, where the basal domain is defined as a minimal regulatory domain surrounding and including each gene, which is defined as 5,000 base pairs upstream to 1,000 base pairs downstream. This basal region is extended by up to 1 megabase, or until it reaches the basal domain of another gene. This means that an ATAC-seq peak can be associated with multiple genes if it is found in the extended regulatory region of both of those genes. Enrichment of specific subsets of the peaks at different distances from nearby genes were calculated using a two-sided hypergeometric test. All tested peaks were used as a background population for each subset of peaks tested.

### Enrichment analysis - ENCODE cis-regulatory elements

ENCODE cis-regulatory element regions were downloaded from https://screen.encodeproject.org/, and a enrichment analysis was carried out much in the same way as described in the previous section. BEDTools was used to identify ATAC-seq peaks that overlapped with ENCODE cis-regulatory elements. A two-sided hypergeometric test was used, with all tested peaks as a null background, to determine if specific differentially accessible ATAC-seq subsets were enriched for the four cis-regulatory element categories - promoter-like, proximal enhancer-like, distal enhancer-like, and ctcf-only elements.

### Enrichment analysis - ChIP-seq datasets

Peaks for 20 transcription factor ChIP-seq experiments in HCT-116 cell line were downloaded from ENCODE, with the exception of the ChIP-seq peaks for the SMARCA4 and SMARCC1 subunits of the SWI/SNF complex which were previously published and made available upon publication (Mathur et al., 2017). Two-sided hypergeometric tests were used as described in the previous methods sections to determine if the differentially accessible peaks were enriched for specific ChIP-Seq signals.

### Enrichment analysis - Kelso et al. Dataset

Publicly available ATAC-seq data from the Kelso et al. study was analyzed using the same pipeline as we used for our own butyrate-treated ATAC-seq data (Fig. 3A). This produced sets of differentially accessible peaks that could then be directly compared to between the two studies. Two-sided hypergeometric tests were used as described in the previous methods sections to determine if there was significant overlap between differentially accessible peaks in the two datasets.

### HOMER motif analysis

The HOMER motif analysis software was used to identify motifs that were enriched in differentially accessible peaks, with all tested peaks used as a background. All *de novo* motifs identified at P < 1e-12 are shown in Fig. 2b. The HOMER findMotifsGenome.pl command was used with the default parameters.

### LD-score regression analysis of colorectal cancer GWAS

Stratified LD score regression was used to test whether colorectal cancer heritability was enriched in peaks that open or close in response to butyrate treatment (Bulik-Sullivan et al., 2015; Finucane et al., 2015, 2018). Using colorectal cancer GWAS data from (Zhou et al., 2018), enrichment was tested in two sets of open (1330 and 3194 peaks respectively) and closed (1453 and 2935 peaks respectively) peaks evaluated against different backgrounds. The regression was adjusted for the set of all background peaks relevant to each enrichment test, and enrichment for open and closed peaks were tested separately. In all tests, we added a 10kb window on either side of each peak.

### Analysis of somatic variants associated with cancer

A total of 828 whole-genome somatic variants VCF files were downloaded from data storage services provided by the Pan-Cancer Analysis of Whole Genomes (PCAWG) project (ICGC/TCGA Pan-Cancer Analysis of Whole Genomes Consortium, 2020). These samples all came from the TCGA wing of the study that was conducted in the USA. Both SNPs and indels were included in the analysis. The somatic mutation rate was calculated in the differentially accessible ATAC-seq peaks for each sample, and then compared to the mutation rate in the non-differentially accessible peaks. This same mutation rate ratio was also calculated for the just the distal peaks. Samples were then grouped by body site of origin, and the log-transformed ratios were tested using a two-sided, one-sample t-test to determine if the regions were enriched or depleted of somatic variations relative to the background mutation rate.

### Cell Counting Assays

Cells were grown in T25 flasks in triplicate - three flasks per condition per treatment. At designated times, cells were trypsinized, resuspended, and counted using a hemocytometer. This was independently repeated three times. Results were visualized using ggplot2 (Wickham, 2016).

### Extracting Nuclear Protein

HCT-116 cells were lysed in 10 mM Tris·Cl, pH 7.4, 10 mM NaCl, 3 mM MgCl2, 0.1% (v/v) Igepal CA-630. The supernatant was removed (cytoplasmic protein). The nuclear pellet was lysed with Radioimmunoprecipitation assay (RIPA) buffer. Protein quantification of nuclear extract was performed using Bicinchoninic acid assay (BCA, Pierce). For total protein extraction to be used for Western blots, however, RIPA buffer (Pierce) was the only lysis buffer utilized.

### HDAC Activity in Nuclear Extracts

HCT-116 nuclear extracts were treated with 1.5 mM butyrate in triplicate. We performed Fluor De Lys HDAC fluorometric activity assay (Enzo Life Sciences) with manufacturer’s protocols, using 6 μg of nuclear extract and 200 μM substrate in a 50 μL total reaction each Fluor De Lys HDAC fluorometric activity assay (Enzo Life Sciences). This was incubated for 2 hours at 37 °C. 50 μL of developer was added. Fluorescence was measured at 350 excitation 450 emission 15 minutes later. This was performed on a Tecan Infinite M1000 Pro. Costar 3628 flat bottom 96 well plates were used for HDAC assays.

### Statistical Analysis of HDAC activity and Cell Number

Overall significance was assessed by One-way ANOVA (ANalysis Of VAriance). Differences between groups were revealed via post-hoc Tukey HSD (Honestly Significant Difference) test. All measurements were visualized as standard error of the mean (SEM).

### Western Blots

25 μg of protein, quantified with BCA, was loaded onto Bolt 10 percent Bis-Tris gels (Invitrogen) and transferred via iBlot 2 technology (Invitrogen). Standard Licor protocol was performed. LI-COR Odyssey Infrared Imaging System was used to visualize results. Western blots were independently validated three times. Anti-Histone H3 (acetyl K9 + K14 + K18 + K23 + K27) antibody (ab47915) was used to measure acetylation of histones. Monoclonal Anti-B-Actin (A5316, Sigma) was used as a control.

## Data and Software Availability

The accession number for the ATAC-Seq and RNA-Seq data generated in this study and reported in this paper can be found in SRA under Bioproject PRJNA715317.

## Supplementary Information

Supplemental Information includes Supplemental Experimental Procedures, six figures, and two tables.

## Acknowledgements

The authors would like to thank Anshul Kundaje and Peyton Greenside for their advice and guidance in running the Big Data Script ATAC-seq pipeline and performing differential analysis. Sequencing costs were supported via NIH S10 Shared Instrumentation Grant (1S10OD02014101), Damon Runyon Clinical Investigator Award to A.S.B., and the Stanford ADRC grant # P50AG047366. A.S.B. was supported by a V Foundation grant, McCormick award, and NIH grants AI148623 and AI143757. B.J.F was supported by the National Science Foundation Graduate Research Fellowship DGE-114747 and by a grant from the Stanford Center for Computational, Evolutionary and Human Genomics. M.D was supported by the National Science Foundation Graduate Research Fellowship. S.B.M. was supported by NIH grants R01AG066490, U01HG009431, R01HL142015 and R01HG008150

## Author Contributions

B.J.F., M.D, S.B.M. and A.S.B conceived of the study. A.S.B and S.B.M supervised the research. B.J.F. generated experimental data. M.D performed bioinformatic analyses. A.R. performed statistical analyses. E.C. initially troubleshot culturing and performing ATAC-Seq on HCT-116 cells. B.J.F and M.D. wrote the manuscript. A.S.B, S.B.M, and E.C. edited the manuscript.

