## Supplemental Information for "Chromatin accessibility changes induced by the microbial metabolite butyrate reveal possible mechanisms of anti-cancer effects"

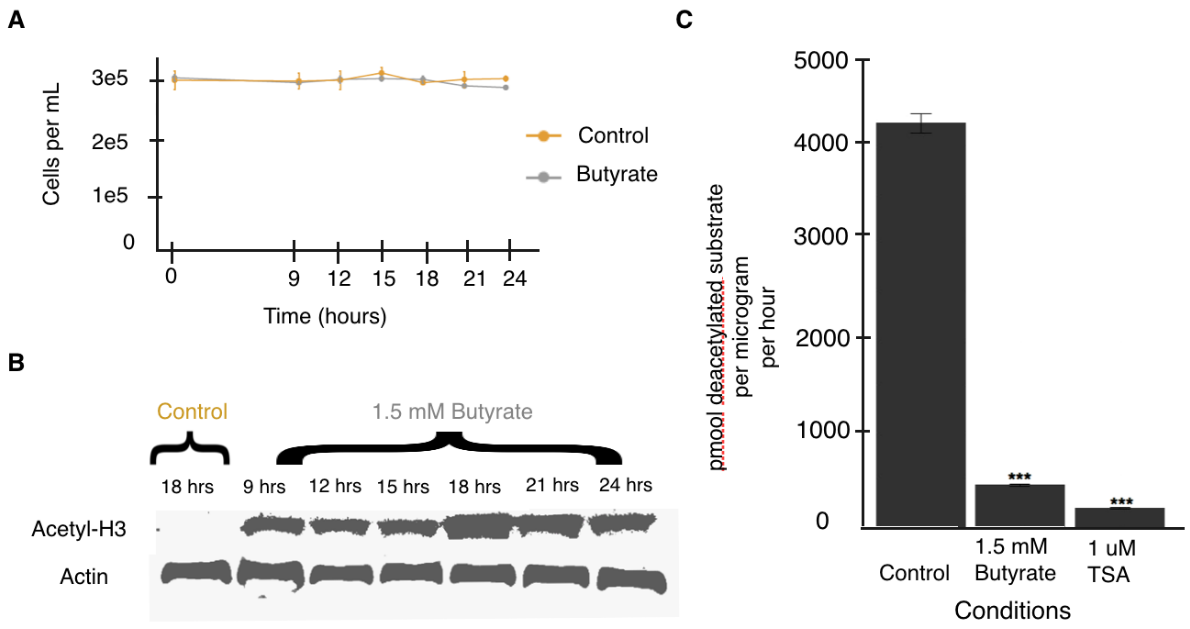


**Figure S1. Butyrate increases histone acetylation and acts as an HDAC inhibitor in HCT-116 cells at 1.5 mM concentrations.** (A) Butyrate at 1.5 mM did not affect cell number and viability over the time course in HCT-116 cells grown in serum-free conditions. (B) Western blot showed butyrate increases histone acetylation in HCT-116 cells at 1.5 mM concentrations. The strongest histone acetylation affect was observed at 18 hours. Actin was used as a control. (C) Butyrate acted as an HDAC inhibitor in HCT-116 cell nuclear extract at 1.5 mM concentrations.

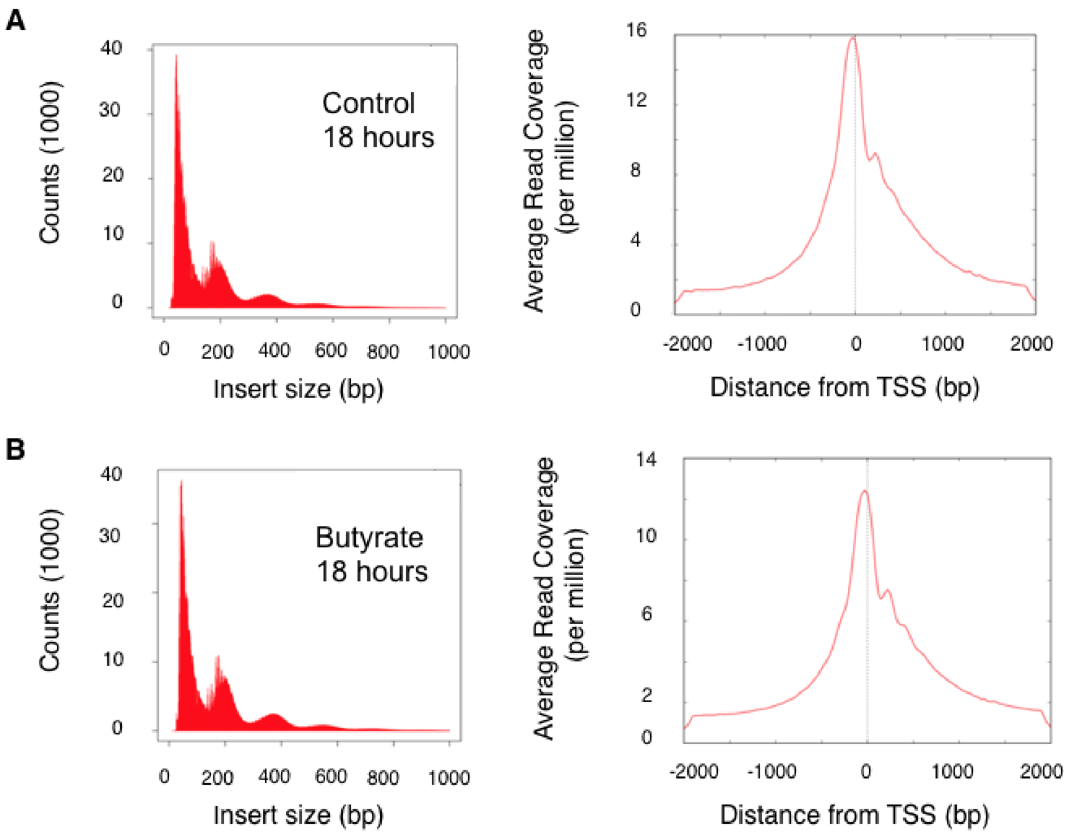


**Figure S2. High-quality ATAC-Seq libraries.** (A) For the control group at 18 hours, we showed pronounced nucleosome phasing patterns and strong enrichment at transcription start sites (TSS), indicative of a high quality ATAC-Seq library. (B) For the butyrate group at 18 hours, we similarly showed that ATAC-seq libraries are of high quality.

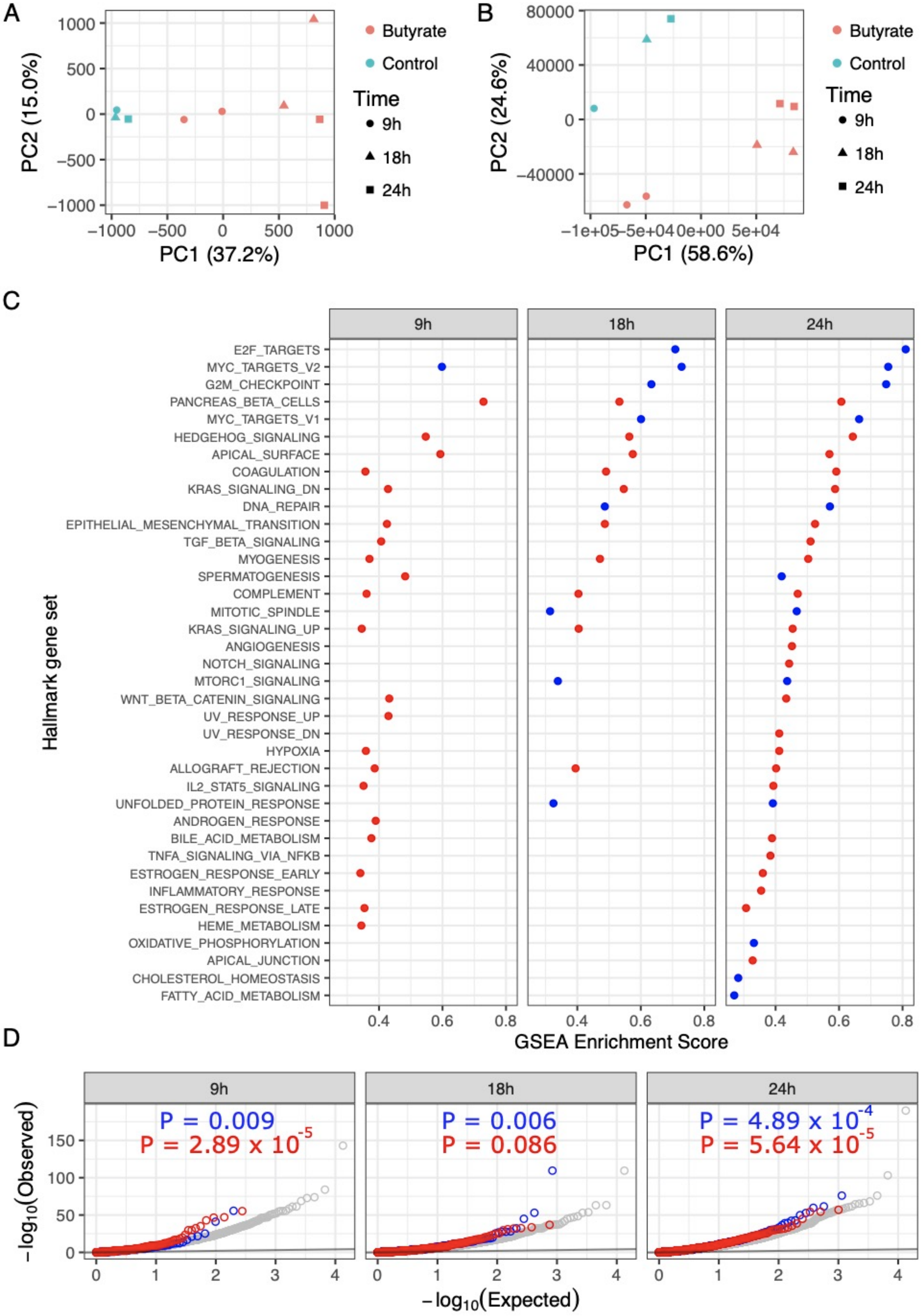
A
PC2 (15.0%)
PC1 (37.2%)
1000
500
0
−500
−1000
−1000
−500
0
500
1000
Butyrate
Control
Time
9h
18h
24h
B
PC2 (24.6%)
PC1 (58.6%)
80000
40000
0
−40000
−1e+05
−5e+04
0e+00
5e+04
Butyrate
Control
Time
9h
18h
24h
C
9h
18h
24h
Hallmark gene set
E2F_TARGETS
MYC_TARGETS_V2
G2M_CHECKPOINT
PANCREAS_BETA_CELLS
MYC_TARGETS_V1
HEDGEHOG_SIGNALING
APICAL_SURFACE
COAGULATION
KRAS_SIGNALING_DN
DNA_REPAIR
EPITHELIAL_MESENCHYMAL_TRANSITION
TGF_BETA_SIGNALING
MYOGENESIS
SPERMATOGENESIS
COMPLEMENT
MITOTIC_SPINDLE
KRAS_SIGNALING_UP
ANGIOGENESIS
NOTCH_SIGNALING
MTORC1_SIGNALING
WNT_BETA_CATENIN_SIGNALING
UV_RESPONSE_UP
UV_RESPONSE_DN
HYPOXIA
ALLOGRAFT_REJECTION
IL2_STAT5_SIGNALING
UNFOLDED_PROTEIN_RESPONSE
ANDROGEN_RESPONSE
BILE_ACID_METABOLISM
TNFA_SIGNALING_VIA_NFKB
ESTROGEN_RESPONSE_EARLY
INFLAMMATORY_RESPONSE
ESTROGEN_RESPONSE_LATE
HEME_METABOLISM
OXIDATIVE_PHOSPHORYLATION
APICAL_JUNCTION
CHOLESTEROL_HOMEOSTASIS
FATTY_ACID_METABOLISM
0.4
0.6
0.8
GSEA Enrichment Score
D
9h
18h
24h
P = 0.009
P = 2.89 x 10^-5
P = 0.006
P = 0.086
P = 4.89 x 10^-4
P = 5.64 x 10^-5
−log10(Observed)
150
100
50
0
0
1
2
3
4
−log10(Expected)

**Figure S3. Additional analysis of RNA-seq and ATAC-seq data.** (A) Principal components analysis of normalized ATAC-seq counts, showing first and second principal components. (B) Principal components analysis of normalized RNA-seq counts, showing first and second principal components. (C) Gene set enrichment analysis (GSEA) results for ranked lists of genes ordered by expression fold change. Showing enrichment scores for hallmark gene sets. Red points indicate gene sets that are enriched among up-regulated genes, and blue points indicate gene sets that are enriched among down-regulated genes. Showing all three time points, and all gene sets that meet a significance cutoff of FDR < 0.25. (D) QQ-plot of p-value statistics of differentially expression analysis. Grey circles indicate all genes, blue circles indicate all genes associated with an ATAC-seq peak that opens in response to butyrate, and red circles indicate all genes associated with an ATAC-seq peak that closes in response to butyrate. P-values indicate the results of two-sided nonparametric Kolmogorov Smirnov tests, where the distribution of p-values in the blue and red gene subsets are compared to the distribution of p-values of all background genes. The black line indicates the null expectation of uniformly-distributed p-values.

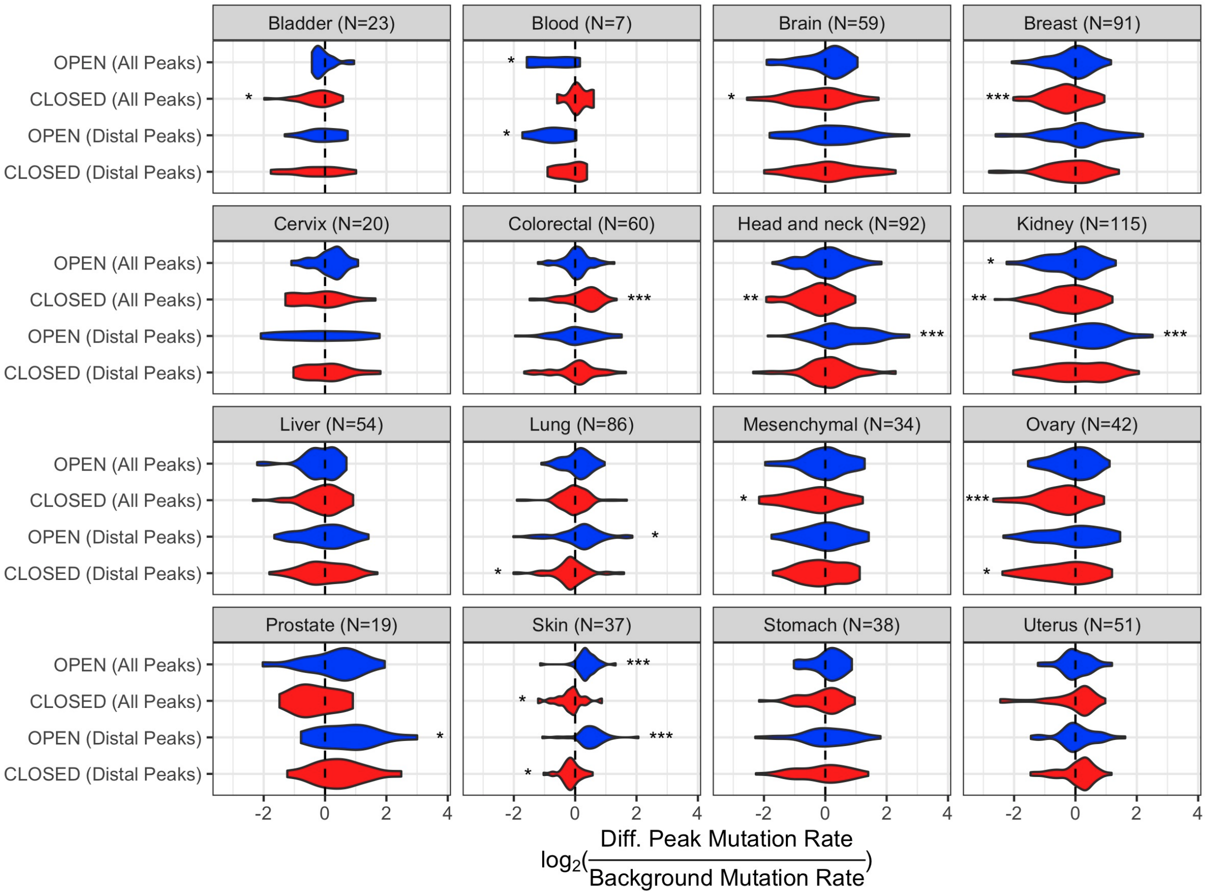


**Figure S4. Somatic variation enrichment within differential ATAC-seq peaks relative to background across tumor body sites.** Showing all 16 cancer body sites included in the downloaded data. Peak sets and colors are the same as in Fig. 4B. Asterisks indicate significance of one-sample t-tests, with ‘*’, ‘**’, and ‘***’ indicating $P < 0.05$, $P < 0.01$, and $P < 0.001$, respectively. The location of the asterisks, either above or below zero along the x-axis, indicate the direction of the enrichment/depletion. Observations were excluded if the mutation rate was too low to estimate a relative mutation rate.
